# Slow oscillation-sleep spindle coupling is associated with fear extinction retention in trauma-exposed individuals

**DOI:** 10.1101/2025.01.27.634866

**Authors:** Dan Denis, Ryan Bottary, Tony J. Cunningham, Per Davidson, Cagri Yuksel, Mohammed R. Milad, Edward F. Pace-Schott

## Abstract

Posttraumatic stress disorder (PTSD) can be characterized as a disorder of fear learning and memory, in which there is a failure to retain memory for the extinction of conditioned fear. Sleep has been implicated in successful extinction retention. The coupling of sleep spindles to slow oscillations (SOs) during non-rapid eye movement sleep has been shown to broadly underpin sleep’s beneficial effect on memory consolidation. However, the role of this oscillatory coupling in the retention of extinction memories is unknown. In a large sample of 124 trauma-exposed individuals, we investigated SO-spindle coupling in relation to fear extinction memory. We found that participants with a PTSD diagnosis, relative to trauma-exposed controls, showed significantly altered SO-spindle timing, such that PTSD participants exhibited spindle coupling further away from the peak of the SO. Across participants, the amount of coupling significantly predicted extinction retention, with coupled spindles uniquely predicting successful extinction retention compared to uncoupled spindles. These results suggest that SO-spindle coupling is critical for successful retention of extinguished fear, and that SO-spindle coupling dynamics are altered in PTSD. These alterations in the mechanics of sleep may have substantial clinical implications, meriting further investigation.

## Introduction

Posttraumatic stress disorder (PTSD) is a condition centered around an emotional declarative (i.e., traumatic) memory in which fear becomes generalized and resistant to extinction (1). PTSD development can be modeled using Pavlovian aversive conditioning paradigms. In these designs, individuals with PTSD, compared to trauma-exposed, non-PTSD controls, show impairments in the retention of learned extinction following a 24-hour delay (2). Extinction learning does not erase pre-existing conditioned fear but rather constitutes a new memory that inhibits a conditioned fear memory when fear-related cues are encountered.

The ability to retain extinction memories is of high clinical relevance, both in terms of underpinning natural resilience to PTSD development following trauma exposure, but also in the therapeutic encoding of extinction during exposure-based therapies (3,4). In healthy participants, post-learning sleep has been demonstrated to play an important role in the consolidation and generalisation of extinction memories (3–8). This process has typically been associated with rapid eye movement (REM) sleep (5–9), though results have not always been consistent (10,11).

Other evidence suggests that non-REM (NREM) sleep oscillations play a mechanistic role in sleep-associated memory consolidation (12,13). Specifically, during periods of NREM, memories are repeatedly reactivated in hippocampal-cortical circuits, leading to overnight memory strengthening. These reactivation processes are governed by the precise temporal coupling of hippocampal sharp-wave ripples (∼80-150Hz), thalamocortical sleep spindles (∼12-15Hz), and global slow oscillations (SOs < 1Hz) (13). Spindles and SOs can be recorded non-invasively in humans using the EEG from standard polysomnography (PSG) sleep studies. Indeed, spindles coupled to SOs are better predictors of overnight consolidation than uncoupled spindles (14–16). Of these coupled spindles, coupling that occurs close to the positive peak of the SO reflects an optimal state for cross-regional dialogue and information transfer between hippocampus and the cortex during sleep (17– 19). Given the established role of the hippocampus in contextualising extinction memory, there is a strong theoretical rationale to hypothesize that SO-spindle mediated hippocampal memory reactivation processes may be critical to the long-term strengthening of extinction memories (20,21).

Trauma memory replay, a process partly dependent on the hippocampus (22), is common in PTSD in the form of intrusive memories and posttraumatic nightmares (23). Contextual fear memory consolidation during sleep in rodents has been linked with the 3-way temporal coupling of hippocampal sharp-wave ripples (indicative of hippocampal replay) with spindles and SOs (24). Recent work has shown that impaired hippocampal responses to threat in the aftermath of a trauma were associated with greater PTSD symptoms at a two-week follow-up (25). Given reciprocal interactions between SO-spindle coupling and hippocampal activity (19), deficits in hippocampal functioning in those with recent trauma exposure may alter coupling patterns in a manner related to their degree of symptomatology. A small but growing number of studies from our group and others have suggested alterations in sleep spindle properties in PTSD (26–30). However, to the best of our knowledge, no studies have examined SO-spindle coupling in trauma exposed patients with and without PTSD.

The aim of this study was to expand our understanding of extinction memory processes in a large sample of trauma-exposed individuals, by testing the hypothesis that there is an association between SO-spindle coupling and extinction recall in this group. These relationships can be directly informative to treatment. Targeted memory reactivation (TMR) paradigms promote spindle-mediated memory reactivation through exposing participants to sound or odor cues presented at learning (12,31). Prior research has demonstrated that externally cueing fear memories during NREM sleep fosters extinction learning (32,33), and TMR has been investigated as a means to augment therapy for PTSD (30). Additionally, SO-spindle coupling can be enhanced via non-invasive stimulation techniques (34). Therefore, SO-spindle coupling may be a marker of extinction memory consolidation during sleep that is amenable to treatment. Here, for the first time, we sought to test the possibility of such relationships in trauma-exposed humans.

## Methods

The data reported here is part of a larger completed study (8,26,35–37). All results presented here are novel and were preregistered on the Open Science Framework (https://osf.io/27nre) unless otherwise noted.

### Participants

One hundred and thirty-three right-handed participants were recruited from the greater Boston metropolitan area using online and posted advertisements. All had experienced a DSM-5 criterion-A traumatic event within the last two years but not within the 30 days prior to study participation. Current and lifetime history of psychiatric disorders were assessed using the Structured Clinical Interview for DSM-IV-TR for Non-Patients (SCID 1/NP; First et al., 2007). PSTD symptoms were assessed using the Clinician-Administered PTSD Scale for DSM-5 (CAPS-5; Weathers et al., 2013) and the PTSD Checklist for DSM-5 (PCL-5). This study followed a Research Domain Criteria (RDoC) design in which dimensional rather than categorical measures were targeted, and PTSD diagnoses were established post-hoc from diagnostic evaluations. A total of 68 participants were diagnosed with PTSD using CAPS-5, and 65 were trauma-exposed controls without PTSD at the time of interview (TEC). Sleep disorders were screened using the Pittsburgh Structured Clinical Interview for Sleep Disorders (SCID-SLD), a widely used (Insana et al. 2013, Stocker et al. 2017) but unpublished, in-house instrument and all participants completed a urine toxicology screen and were negative for 11 abused substances. An upper age limit of 40 was imposed to ensure participants retained a measurable amount of slow wave sleep. Study procedures were reviewed and approved by the Massachusetts General Hospital Institutional Review Board. Participants provided written informed consent and were paid for their participation.

For the present analyses, a total of 124 participants (PTSD: N=62, TEC: N=62) provided usable data. A smaller number (N=87; PTSD: n=41, TEC: n=46) was used for correlations with skin conductance responses (SCRs) after removing SCR non-responders (see below). Demographic and clinical characteristics of the sample used for analyses here are presented in **Table 1**.

**Table 1.**
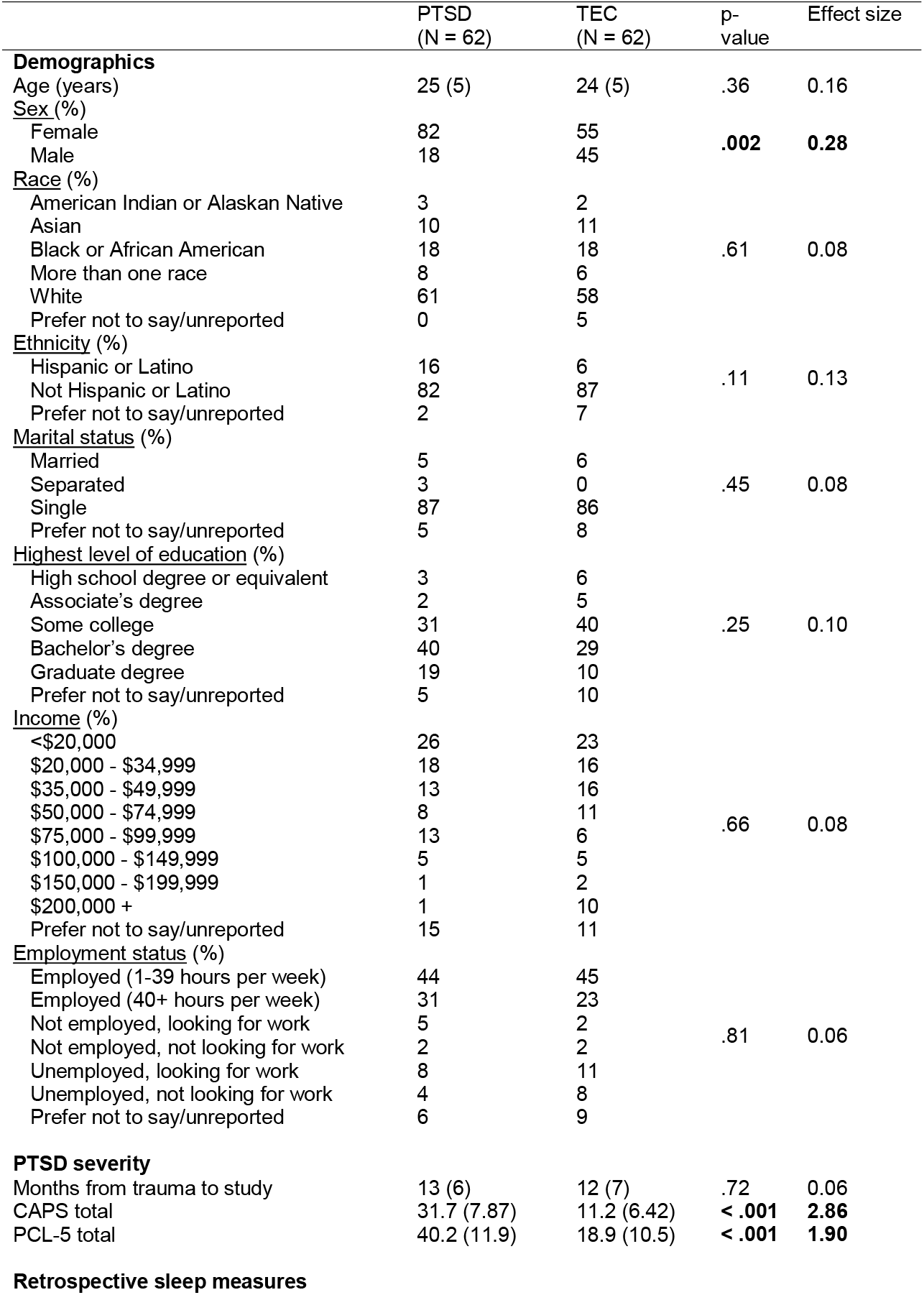
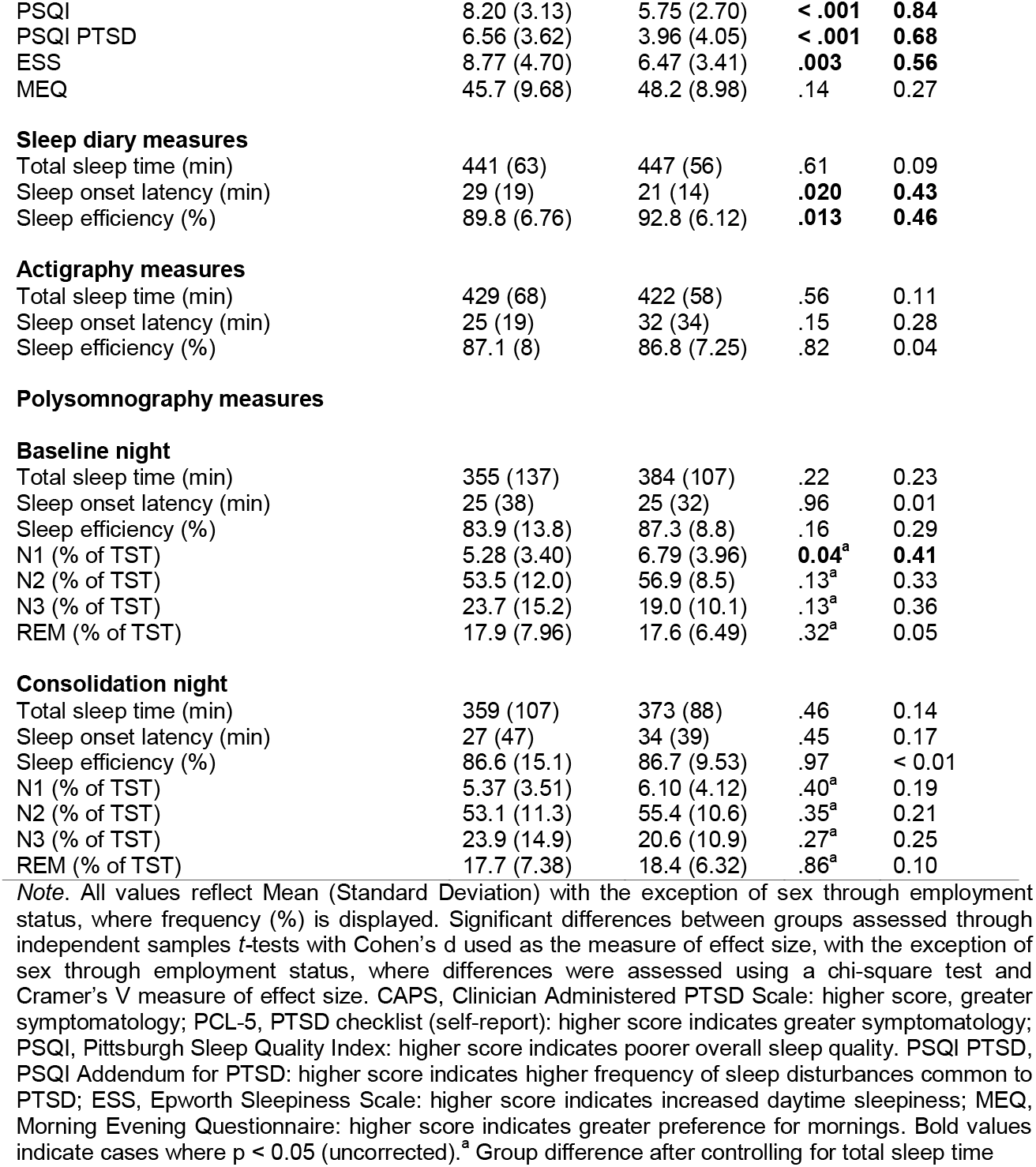
Sample characteristics.

|  | PTSD<br>(N = 62) | TEC<br>(N = 62) | p-value | Effect size |
| --- | --- | --- | --- | --- |
| **Demographics** |  |  |  |  |
| Age (years) | 25 (5) | 24 (5) | .36 | 0.16 |
| <u>Sex (%)</u> |  |  |  |  |
| Female | 82 | 55 | **.002** | **0.28** |
| Male | 18 | 45 |  |  |
| <u>Race (%)</u> |  |  |  |  |
| American Indian or Alaskan Native | 3 | 2 | .61 | 0.08 |
| Asian | 10 | 11 |  |  |
| Black or African American | 18 | 18 |  |  |
| More than one race | 8 | 6 |  |  |
| White | 61 | 58 |  |  |
| Prefer not to say/unreported | 0 | 5 |  |  |
| <u>Ethnicity (%)</u> |  |  |  |  |
| Hispanic or Latino | 16 | 6 | .11 | 0.13 |
| Not Hispanic or Latino | 82 | 87 |  |  |
| Prefer not to say/unreported | 2 | 7 |  |  |
| <u>Marital status (%)</u> |  |  |  |  |
| Married | 5 | 6 | .45 | 0.08 |
| Separated | 3 | 0 |  |  |
| Single | 87 | 86 |  |  |
| Prefer not to say/unreported | 5 | 8 |  |  |
| <u>Highest level of education (%)</u> |  |  |  |  |
| High school degree or equivalent | 3 | 6 | .25 | 0.10 |
| Associate's degree | 2 | 5 |  |  |
| Some college | 31 | 40 |  |  |
| Bachelor's degree | 40 | 29 |  |  |
| Graduate degree | 19 | 10 |  |  |
| Prefer not to say/unreported | 5 | 10 |  |  |
| <u>Income (%)</u> |  |  |  |  |
| <\$20,000 | 26 | 23 | .66 | 0.08 |
| \$20,000 - \$34,999 | 18 | 16 | | |
| \$35,000 - \$49,999 | 13 | 16 | | |
| \$50,000 - \$74,999 | 8 | 11 | | |
| \$75,000 - \$99,999 | 13 | 6 | | |
| \$100,000 - \$149,999 | 5 | 5 | | |
| \$150,000 - \$199,999 | 1 | 2 | | |
| \$200,000 + | 1 | 10 | | |
| Prefer not to say/unreported | 15 | 11 |  |  |
| <u>Employment status (%)</u> |  |  |  |  |
| Employed (1-39 hours per week) | 44 | 45 | .81 | 0.06 |
| Employed (40+ hours per week) | 31 | 23 |  |  |
| Not employed, looking for work | 5 | 2 |  |  |
| Not employed, not looking for work | 2 | 2 |  |  |
| Unemployed, looking for work | 8 | 11 |  |  |
| Unemployed, not looking for work | 4 | 8 |  |  |
| Prefer not to say/unreported | 6 | 9 |  |  |
| **PTSD severity** |  |  |  |  |
| Months from trauma to study | 13 (6) | 12 (7) | .72 | 0.06 |
| CAPS total | 31.7 (7.87) | 11.2 (6.42) | **< .001** | **2.86** |
| PCL-5 total | 40.2 (11.9) | 18.9 (10.5) | **< .001** | **1.90** |
| Retrospective sleep measures |  |  |  |  |
| PSQI | 8.20 (3.13) | 5.75 (2.70) | **< .001** | **0.84** |
| PSQI PTSD | 6.56 (3.62) | 3.96 (4.05) | **< .001** | **0.68** |
| ESS | 8.77 (4.70) | 6.47 (3.41) | **.003** | **0.56** |
| MEQ | 45.7 (9.68) | 48.2 (8.98) | .14 | 0.27 |
| **Sleep diary measures** |  |  |  |  |
| Total sleep time (min) | 441 (63) | 447 (56) | .61 | 0.09 |
| Sleep onset latency (min) | 29 (19) | 21 (14) | **.020** | **0.43** |
| Sleep efficiency (%) | 89.8 (6.76) | 92.8 (6.12) | **.013** | **0.46** |
| **Actigraphy measures** |  |  |  |  |
| Total sleep time (min) | 429 (68) | 422 (58) | .56 | 0.11 |
| Sleep onset latency (min) | 25 (19) | 32 (34) | .15 | 0.28 |
| Sleep efficiency (%) | 87.1 (8) | 86.8 (7.25) | .82 | 0.04 |
| **Baseline night** |  |  |  |  |
| Total sleep time (min) | 355 (137) | 384 (107) | .22 | 0.23 |
| Sleep onset latency (min) | 25 (38) | 25 (32) | .96 | 0.01 |
| Sleep efficiency (%) | 83.9 (13.8) | 87.3 (8.8) | .16 | 0.29 |
| N1 (% of TST) | 5.28 (3.40) | 6.79 (3.96) | **0.04<sup>a</sup>** | **0.41** |
| N2 (% of TST) | 53.5 (12.0) | 56.9 (8.5) | .13<sup>a</sup> | 0.33 |
| N3 (% of TST) | 23.7 (15.2) | 19.0 (10.1) | .13<sup>a</sup> | 0.36 |
| REM (% of TST) | 17.9 (7.96) | 17.6 (6.49) | .32<sup>a</sup> | 0.05 |
| **Consolidation night** |  |  |  |  |
| Total sleep time (min) | 359 (107) | 373 (88) | .46 | 0.14 |
| Sleep onset latency (min) | 27 (47) | 34 (39) | .45 | 0.17 |
| Sleep efficiency (%) | 86.6 (15.1) | 86.7 (9.53) | .97 | < 0.01 |
| N1 (% of TST) | 5.37 (3.51) | 6.10 (4.12) | .40<sup>a</sup> | 0.19 |
| N2 (% of TST) | 53.1 (11.3) | 55.4 (10.6) | .35<sup>a</sup> | 0.21 |
| N3 (% of TST) | 23.9 (14.9) | 20.6 (10.9) | .27<sup>a</sup> | 0.25 |
| REM (% of TST) | 17.7 (7.38) | 18.4 (6.32) | .86<sup>a</sup> | 0.10 |

*Note.* All values reflect Mean (Standard Deviation) with the exception of sex through employment status, where frequency (%) is displayed. Significant differences between groups assessed through independent samples *t*-tests with Cohen's *d* used as the measure of effect size, with the exception of sex through employment status, where differences were assessed using a chi-square test and Cramer's *V* measure of effect size. CAPS, Clinician Administered PTSD Scale: higher score, greater symptomatology; PCL-5, PTSD checklist (self-report): higher score indicates greater symptomatology; PSQI, Pittsburgh Sleep Quality Index: higher score indicates poorer overall sleep quality. PSQI PTSD, PSQI Addendum for PTSD: higher score indicates higher frequency of sleep disturbances common to PTSD; ESS, Epworth Sleepiness Scale: higher score indicates increased daytime sleepiness; MEQ, Morning Evening Questionnaire: higher score indicates greater preference for mornings. Bold values indicate cases where $p < 0.05$ (uncorrected).<sup>a</sup> Group difference after controlling for total sleep time

### Sleep Assessment and PSG Monitoring

Following psychological screening, participants completed a ∼2-week sleep-assessment period (mean 15±3 days) during which they continuously wore an Actiwatch-2 device (Phillips Respironics, Bend, OR) and filled out daily sleep and nightmare dairies. Over this period, participants also completed an online battery of questionnaires assessing trauma history, habitual sleep, circadian preference, anxiety, mood and personality variables using the Research Electronic Data Capture system (40,41). Approximately midway through the 14-day period, participants completed a combined sleep disorders diagnostic and acclimation night using ambulatory polysomnography (PSG). This was followed by a second PSG recording (“Baseline night”). The following evening, participants then completed the Fear Conditioning and Extinction Learning phases of a validated fear-conditioning and extinction protocol (21,42), during fMRI scanning (37). Following fear conditioning and extinction learning, participants completed a third PSG recording (“Consolidation night”). The following day, 24-hr after fear conditioning and extinction, participants returned to the lab for the Extinction Recall phase of the Pavlovian conditioning and extinction task, again conducted during fMRI scanning. Alcohol and recreational drugs were prohibited throughout the protocol and, beginning on the day before the Baseline Night, caffeine use and daytime naps were also prohibited.

Sleep was monitored using the Somte-PSG device (Compumedics USA, Inc., Charlotte, NC). Electrodes were attached in the laboratory, then participants were sent home to sleep. The montage included six EEG channels (F4, F4, C3, C4, O1, O2) referenced to contralateral mastoids, two electrooculography channels, two submental electromyography channels, and two electrocardiography channels (positioned below the right clavicle and on the left 5^th^ intercostal space). The first sleep recording was considered an acclimation and sleep-disorders screening night and included additional channels for respiration transducer belts, pulse-oximeter, and tibialis movement. Signals were recorded at 256Hz. All PSG nights were scored by an experienced, RSPGT-certified research polysomnographer following the American Academy of Sleep Medicine criteria (43). The acclimation/assessment night was additionally examined by the same scorer for clinically significant obstructive sleep apnoea (OSA) and periodic leg movement disorder (PLMD). No participants were excluded on these grounds.

### Pavlovian conditioning task

A validated 2-day paradigm was used to probe fear conditioning and extinction during ongoing fMRI recording (46,47; **Figure 1**). The task used in the present investigation had four phases: Habituation, Fear Conditioning, Extinction Learning and Extinction Recall. During Habituation, participants were presented with all combinations of to-be-conditioned stimuli (three separate colored lamps) in each of the study contexts (Conditioning context, Extinction context). During trials in all phases, the contexts first appeared for 3 sec after which the colored-light conditioned stimuli (CS) appeared for an additional 6 sec. Stimuli were projected onto a viewing screen inside of the MRI scanner. During Fear Conditioning, participants were conditioned to two colored lamps (CS+1 and CS+2; colors counterbalanced across participants), but not a third (CS-), using a mild finger shock delivered at stimulus offset (i.e., when the study context and lamp disappeared from the computer screen). During Extinction Learning, which immediately followed fear conditioning, one previously conditioned colored lamp (CS+E), but not the other (CS+U), was extinguished by repeatedly presenting the stimulus without shock reinforcement in the extinction context. Presentation of the CS-were interspersed throughout extinction learning. During Extinction Recall, which took place the following day and in the extinction context, all three lamps were presented without shock reinforcement (for additional details, see (44)).

**Figure 1.**
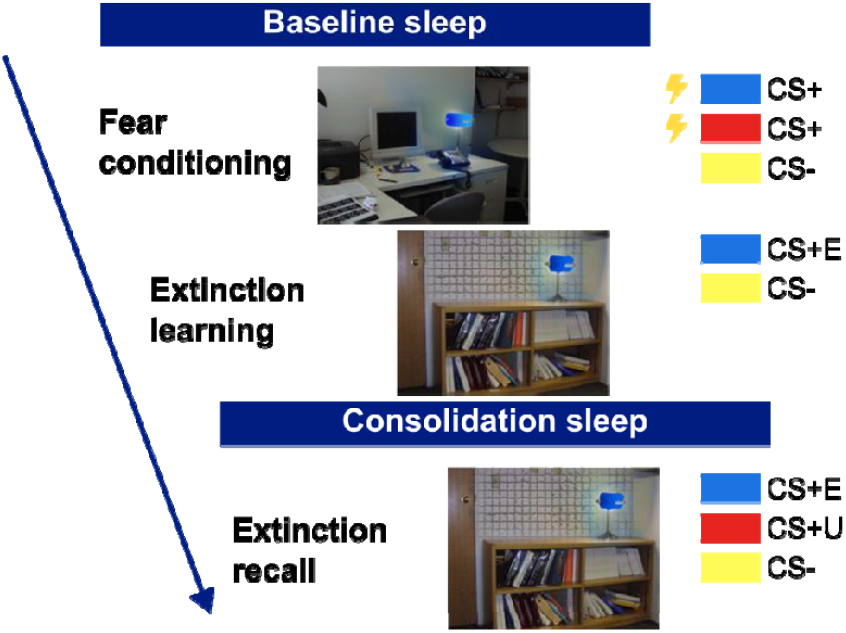
Pavlovian fear conditioning task. Following a baseline night of sleep, participants were conditioned to associate two different colored lights with a mild electric shock (Fear Conditioning). A third stimulus was never paired with a shock. Immediately after this, participants underwent extinction learning where one of the previously conditioning stimuli was represented with no electric shocks. 24 hours later (including a second night of sleep), participants underwent extinction recall where all three stimuli were represented without any shocks being administered.

Immediately following each phase except Habituation, participants verbally reported shock expectancy for the first and last presentations of each CS appearing during that phase on a scale from 1 (“not expecting a shock at all”) to 5 (“expecting a shock very much”).

### Skin Conductance Monitoring and Electric Shocks

Skin conductance levels of participants’ left hand were continuously monitored throughout each study phase at 200Hz using a BIOPAC MP150 system with Acqknowledge 4.3 software (BIOPAC Systems, Inc., Goleta, CA). Stimulating electrodes attached to participants right index and middle finger delivered mild electric shocks. Finger shock level was calibrated for each participant by administering increasing intensities of shock in 5 increments (0.8 – 4 mA) using the Coulbourn Transcutaneous Aversive Finger Stimulator (Coulbourn Instruments, Allentown, PA). Participants selected a level of shock that they perceived to be “highly annoying but not painful”. Anticipatory skin conductance responses (SCRs) were used to measure reactivity to all study stimuli (see **Supplementary Materials**).

### Extinction recall and generalisation indices

Measures of extinction recall and generalisation were calculated for the purposes of correlating with sleep physiology. For subjective shock expectancy ratings, a subjective extinction retention index (sERI) was calculated as the expectancy to the first CS+E trial during Extinction recall divided by the mean expectancy for the last two CS+s during Fear conditioning, yielding a percentage of the maximal conditioned fear that had been retained (6,44). A lower sERI reflects better extinction retention. A subjective extinction generalisation index (sEGI) was calculated as the expectancy to the first CS+U at Extinction recall divided by the mean expectancy of the last two of each CS+ during Fearing conditioning minus the sERI (45). This yields a percentage of the maximum conditioned fear represented by the retained difference in response to an unextinguished vs extinguished CS+. These indices were selected to align with previous publications from our group, acknowledging that different extinction indices may not intercorrelate (46). Analogous measures were calculated for SCRs (see **Supplementary Materials**).

### Slow oscillation-spindle coupling

Sleep spindles were detected during NREM sleep using an automated wavelet-based detector. To capitalize on the large individual differences in spindle peak frequency (47), we tuned the detector around each participant’s most prominent sigma band peak. Spindle peak frequency was identified through visual inspection of the NREM power spectrum, calculated as the power spectral density (PSD) for NREM sleep at central (C3, C4) electrodes (9,26). PSD was estimated on the derivative of the time series to minimize 1/*f* scaling using Welch’s method with five second windows and 50% overlap (26,47–49). The largest, most prominent peak between 12.5 – 16Hz was retained as that participant’s spindle peak frequency. We focused on this frequency range for the “fast” spindle subtype (12.5 – 16Hz) due to their theoretical relevance to mechanisms of sleep-based memory consolidation (12,13). In cases where no discernible peak could be identified, peak frequency was set to 14Hz (50,51).

After deriving each participant’s individualized peak, spindles were automatically detected using a validated detector (14,15,52). The raw EEG signal was subjected to a time-frequency decomposition using complex Morlet wavelets, with the peak frequency of the wavelet set to the individual’s spindle peak and a bandwidth (full width half-max) of 1.3Hz centred on the peak frequency. Spindle detection was performed on the squared wavelet coefficients after being smoothed with a 100ms moving average. A spindle was detected whenever the wavelet signal exceeded a threshold of six times the median signal amplitude of NREM sleep for at least 0.4s. The threshold of 6 times the median has been empirically determined to maximize between class (“spindle”, “nonspindle”) variance (14,51).

Slow oscillations (SOs) were detected using a second automated algorithm (14,15). First, data were bandpass filtered between .5-4Hz and all positive-to-negative zero crossings identified. Candidate slow oscillations were marked if two consecutive crossings fell 0.8 – 2 seconds apart, i.e. only oscillations with a frequency of .5 – 1.25Hz were considered. Oscillations in the top quartile of peak-to-peak amplitudes (i.e., the largest) were retained as slow oscillations (14,53,54).

SO-spindle coupling was identified using a co-occurrence approach. EEG data were bandpass filtered in the individualized spindle band (i.e., the peak frequency with a 1.3Hz bandwidth). Then, the Hilbert transform was applied to extract the instantaneous phase of the delta (.5-4Hz) filtered signal and the instantaneous amplitude of the spindle filtered signal. For each detected spindle, the peak amplitude was determined. Next, it was determined whether the spindle peak occurred within the time course of any detected SOs. For all identified co-occurrences, the phase angle of the SO at the spindle peak was determined. For each participant, we extracted the % of spindles coupled to a slow oscillation, the mean coupling phase (degrees), and the coupling consistency (mean vector length), and averaged these parameters across the two central electrodes.

### Statistical analysis

#### Primary analyses

Group differences in shock expectancy ratings were assessed with a linear mixed effects model with the factors Group (TEC, PTSD), Stimulus type (either CS+E, CS-for extinction recall, or CS+E, CS+U for extinction generalisation) and their interaction entered as fixed effects, and participant entered as a random effect.

Differences in the amount and consistency of SO-spindle coupling between groups was assessed via two linear mixed effects models with the factors Group (TEC, PTSD), Night (baseline, consolidation), and their interaction entered as fixed effects. Participant was entered as a random effect. For coupling phase we first determined whether, across participants, the distribution of coupling phases was non-uniform (i.e., was there a preferential phase for the coupling of spindles to SOs). This was tested using Rayleigh’s test of non-uniformity. To test for the effect of Group and Night on coupling phase, we utilised a 2 (Group: TEC, PTSD) X 2 (Night: baseline, consolidation) ANOVA for circular data (55). A *p* value of < .017 (i.e. 0.05 / 3 comparisons) was deemed statistically significant.

To assess the associations between SO-spindle coupling and extinction measures, sERI/sEGI was regressed against either % coupled spindles or coupling consistency, group (TEC, PTSD), night (Baseline, Consolidation), and their interaction using robust linear regression models. Associations between sERI/sEGI and coupling phase were assessed using circular-linear correlations, separately for each group. This led to a total of eight analyses of extinction and SO-spindle coupling being performed, thus *p* values were adjusted for eight comparisons (i.e., 0.05 / 8 comparisons). Therefore, a *p* value of < .006 was deemed statistically significant.

#### Secondary analyses

Consistent with our previous report, we also repeated the primary analysis of group differences instead comparing participants with low or high levels of symptomatology, defined by the total CAPS score (26). We defined low symptomatology participants as those with a total CAPS score of ten or less (N = 32), and high symptomatology participants as those with a total CAPS score of more than 22 (N = 61). We also reported correlations between total CAPS score and our measures of interest. These additional analyses are more in keeping with the Research Domain Criteria (RDoC) design of this study, in which dimensional rather than categorical measures were targeted. These analyses were not preregistered.

#### Exploratory analyses

For associations between sleep and extinction measures, we focused on subjective measures of extinction recall rather than SCRs due to theoretical (a hippocampal component that may be more influenced by SO spindle coupling (54,56) and practical (a larger sample size resulting in more statistical power, lesser general habituation) considerations. However, for full completeness we report all associations between sleep parameters and SCR measures of extinction in the **Supplementary materials**. No group differences or associations with SO-spindle coupling were observed.

## Results

### Behaviour

Shock expectancy on the first recall trial was significantly higher for the CS+E compared with the CS-(*F*(1,226)=24.38, *p*<.001), with no difference between the CS+E compared with the CS+U (*F*(1,223) = 1.08, *p*=.30); **Figure S1**. There was no main effect of Group on shock expectancy, nor did group interact with Stimulus type (*p*s>.41). Neither sERI or sEGI (see **Methods**) differed between TEC and PTSD groups (all *p*s>.55), between low and high symptomatology subgroups (all *p*s>.88) or correlated with total CAPS score (*r*s<.07, *p*s>.52).

### Group differences in slow oscillations-spindle coupling

A summary of spindle and SO measures is provided in **Table 2**. Across all participants, 16% of spindles were coupled to a slow oscillation (baseline night: 15.76%±4.05%; consolidation night: 15.81%±4.13%), in line with previous findings (14,57). There was evidence of non-uniformity in the preferred phase of the SO to which spindles were coupled (baseline night: Z=81.47, *p*<.001; consolidation night: Z=96.13, *p*<.001). Spindles preferentially coupled just after the peak of the SO (baseline night: 25.82°±24.69°; consolidation night: 24.79°±21.98°). There was a significant main effect of group on coupling phase (F (1,170)=19.50, *p*<.001; **Figure 2A**). On average, the spindles of those participants with a PTSD diagnosis coupled later (33.01°±11.53°) in the SO phase (i.e., further away from the peak) compared to TEC (19.26°±24.75°). Coupling phase did not differ across nights (*F*(1,170)=0.27, *p*=.60), and there was no group ^*^ night interaction (*F*(1,170)=0.40, *p*=.53).

**Table 2.** slow oscillation-spindle coupling metrics.

|  | PTSD<br>M (SD) | TEC<br>M (SD) | p-value | Effect<br>size |
| --- | --- | --- | --- | --- |
| Coupled spindle density (#/min) | 1.03 (0.35) | 0.99 (0.33) | .59 | 0.12 |
| Uncoupled spindle density (#/min) | 5.52 (1.48) | 5.35 (1.37) | .57 | 0.12 |
| % spindles coupled | 16.47 (4.20) | 15.38 (3.50) | .64 | 0.10 |
| % SOs coupled | 16.88 (5.03) | 16.21 (5.68) | .56 | 0.12 |
| Coupling phase (°) | 33.01 (11.53) | 19.26 (24.75) | **< .001** | **0.65** |
| Coupling consistency (vector length) | .41 (.13) | .41 (.14) | .88 | 0.03 |

Note. M = mean, SD = standard deviation. Coupling phase assessed using the circular mean and standard deviation. 0° represents that the peak of the spindle occurred at the positive peak of the slow oscillation. All values reflect the average of the two nights, and the average of electrodes C3 and C4. p-value is for the main effect of group assessed via independent samples t-test (Watson-Williams test in the case of Coupling phase), uncorrected for multiple comparisons. Effect sizes are Cohen's d.

**Figure 2.**
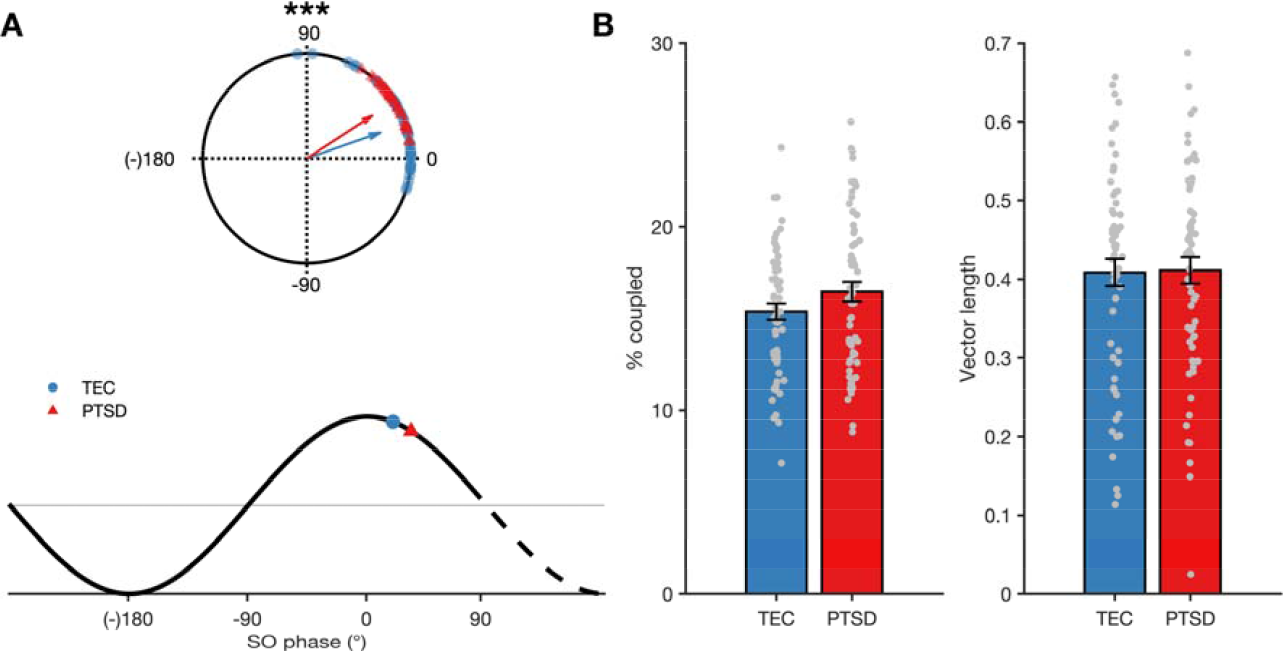
SO-spindle coupling. A – Top: Group differences in spindle coupling phase. Bottom: Mapping of circular phase plots onto slow oscillation waveform. A coupling phase of 0° corresponds to the peak of the spindle occurring at the positive peak of the SO. B – Group differences in overall amount of coupling (left) and consistency of SO-spindle timing (right). Error bars indicate standard error.

The same pattern of results emerged when we compared coupling phase between the low (15.63°±20.40°) and high symptomatology (33.80°±12.75°) subgroups (*F*(1,60)=17.55, *p*<.001), and when total CAPS score was correlated with mean coupling phase (*r*=.33, *p*=.001). Together, this pattern of results suggests that a PTSD diagnosis was associated with a mean coupling phase that is further away from the peak of the slow oscillation. Such effects were restricted to the mean coupling phase. There were no group differences in either the overall amount of coupling or the consistency of coupling timing (*ps* >.10, **Figure 2B**). Neither amount or consistency of coupling differed between high and low symptomatology participants (*ps*>.26) and neither correlated with total CAPS score (*p*s>.19).

### Effect of SO-spindle coupling on extinction retention and generalisation

There was a significant relationship between the overall amount of SO-spindle coupling and sERI (*F*(1,188)=8.83, *p*=.003, *r*=-.23; **Figure 3**). Participants who had a higher percentage of spindles coupled to SOs had a lower sERI value, indicative of better extinction retention as a function of an increased proportion of coupled spindles. There were no interactions with group or night, and neither coupling phase nor consistency predicted sERI (*p*s>.07), suggesting the amount of coupling to uniquely predict extinction retention across all participants.

**Figure 3.**
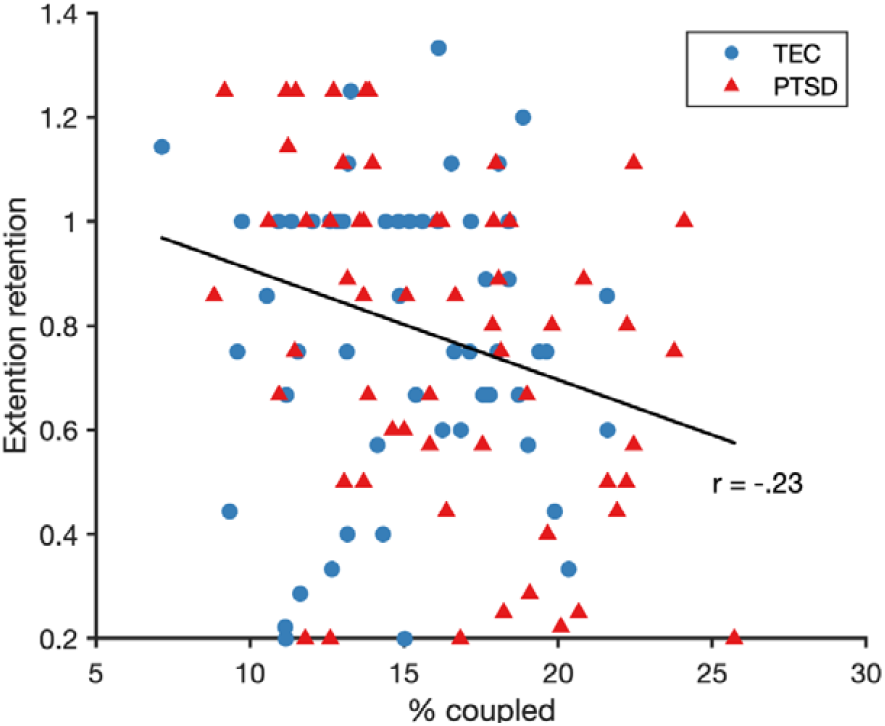
A higher amount of SO-spindle coupling (% spindles coupled to an SO) predicted better subjective (shock expectancy) extinction retention across all participants. A lower extinction retention score indicates better extinction retention. The trend line depicts the robust linear fit.

To further confirm the role of coupling in extinction retention, two supplementary (and un-preregistered) analyses were performed. First, to confirm the importance of SO-spindle coupling in successful extinction, we reasoned that the number of coupled spindles should uniquely predict retention independently of uncoupled spindles (i.e., spindles that did not occur during an SO). To test this, we ran a multiple linear regression predicting sERI from the coupled and uncoupled spindle density (number/min of coupled and uncoupled spindles). The overall regression model was significant (F(1,114)=4.54, p=.013, adjusted R^2^=.06). Critically, only the coupled spindle density predicted sERI (Estimate[95% CI] = - 0.31[-0.52, -0.10], p=.004). Uncoupled spindle density did not predict sERI independently of coupled spindle density (Estimate[95% CI]=0.05[-0.01, 0.10], p=.06).

Second, our preregistered outcome measure of the amount of coupling was the % of spindles that coupled to an SO. However, it is also possible to calculate the % of SOs coupled to a spindle. Whilst the former measure emphasises proportion of spindles that are coupled, that latter emphasises the proportion of coupled SOs. Therefore, to confirm whether the relationship between SO-spindle coupling and extinction retention was primarily driven by spindles or SOs, we ran a second multiple robust linear regression predicting sERI from the % of spindles coupled to an SO and the % of SOs coupled to a spindle as independent predictors. The overall regression model was significant (F(1,114)=3.84, p=.024, adjusted R^2^=.05), with only the % of spindles coupled to an SO predicting sERI (Estimate[95% CI] =-0.02[-0.04, -0.01], p=.017). The % of SOs coupled to a spindle did not independently predict sERI (Estimate[95% CI]=0.001[-0.01, 0.01], p=.92). Thus, this analysis further emphasizes the importance of sleep spindles in promoting extinction retention.

## Discussion

There has been growing interest in the role of non-rapid eye movement (NREM) sleep oscillations in the pathophysiology of PTSD (27). Here, we investigated SO-spindle coupling in PTSD and how it relates to the consolidation of fear extinction memories. We found evidence of altered SO-spindle timing in PTSD. Across all participants, higher amounts of SO-spindle coupling were positively associated with the successful retention of subjective extinction memory. We discuss each of our findings in greater detail below.

We found that the spindles of PTSD patients coupled later in the SO cycle than controls, meaning that spindle coupling occurred further away from the SO peak in participants with PTSD. The coupling of SOs and spindles facilitates information transfer and cross regional dialogue between hippocampus and neocortex (19), with optimal cortical-subcortical interactions occurring when spindles couple to the positive peak of the SO (17,19). The finding that individuals with PTSD show coupling further away temporally from the SO peak points towards an impairment in hippocampal-neocortical dialogue during sleep in PTSD. This finding is consistent with studies of resting-state functional connectivity in PTSD, that consistently find aberrant connectivity between hippocampus and areas of prefrontal (PFC) and posterior cingulate cortices in PTSD compared to controls (58–60). Notably, PFC is a key generator site for SOs during sleep (61,62). Therefore, if reduced hippocampus-PFC connectivity during wake extends into sleep, this could impede cross-regional dialogue and information transfer during sleep that is believed to be mediated by SO-spindle coupling (17,19,63).

Although spindles have been shown to support hippocampal-cortical connectivity during sleep, spindles are initially generated by the reciprocal interactions of thalamocortical neurons (64). Recent work has also implicated the thalamus in the organisation of SOs in the neocortex (65). Therefore, our finding of altered SO-spindle coupling dynamics could also point to an abnormality in thalamic functioning in PTSD. This hypothesis aligns with other strands of research. Sensory gating, one of the primary functions of the thalamus, is impaired in PTSD (66) and has been proposed to contribute to the recurrence of intrusive memories characteristic of the disorder (67). PTSD patients also show reduced thalamic volume (68,69) and alterations in thalamocortical connectivity relative to controls (70,71). Interestingly, spindle density has been shown to correlate with thalamocortical connectivity in both healthy participants and patients with schizophrenia (72). As such, future studies that are able to link resting state functional connectivity to SO-spindle coupling will be immensely valuable in further our understanding of mechanisms underlying altered SO-spindle coupling in PTSD.

Despite this group difference, spindle coupling phase was not predictive of fear extinction retention. Instead, across all participants, the overall amount of coupling correlated with successful subjective extinction retention. SO-spindle coupling has emerged as a candidate mechanism supporting systems-level consolidation during sleep (13). Under this framework, memories are reactivated within SO-spindle complexes, promoting their long-term strengthening (12,73,74). Our finding that more frequent coupling events were associated with greater retention of extinction memory could be interpreted to mean that more frequent opportunities for reactivation of the extinction memory during sleep led to a stronger representation during the recall session. Supporting this, research using TMR to externally reactivate memories during sleep by replaying sound or odor cues presented during learning (75) has found that re-exposing participants during SWS to an odor associated with the CS+ fosters new extinction learning (32,33). The present study complements this prior work by linking the neural oscillations responsible for endogenous memory reactivation (SO-spindle events) to the consolidation of extinction memories in a clinically relevant sample of trauma-exposed individuals.

We observed no effect of experimental night on spindle coupling measures, and the association between coupling and extinction retention did not interact with experimental night. Our result is consistent with other studies showing these oscillations to be highly stable across nights (47,76), suggesting them to be trait-like with a strong genetic component (77). Under this interpretation, our results suggest that a later spindle-coupling phase is a trait-like feature of the sleep EEG that may put one at a heightened risk of developing PTSD following a trauma. Similarly, a higher amount of SO-spindle coupling could also be a trait-like feature that facilitates generally enhanced memory consolidation, including for extinction memories. Alternatively, delayed SO-spindle coupling could also have resulted from certain individual’s reaction to trauma and have developed alongside PTSD. Clearly, prospective studies with longitudinal designs are required to test these hypotheses.

A limitation of the work is that all our analyses were correlational in nature, and we cannot definitively equate more SO-spindle coupling to increased reactivation of the extinction memory. Recent work using multivariate classification approaches have been able to detect a reinstatement of memory content within SO-spindle complexes (74). Additionally, targeted memory reactivation (TMR) paradigms have demonstrated that externally cueing an odor presented during fear conditioning during sleep enhances extinction learning, adding more causal evidence for a role of memory reactivation in fear extinction processes (32). Applying these experimental paradigms and analysis techniques in trauma-exposed participants would be an interesting avenue for future research. Previously, it has been suggested that sleep may affect the balance of strength between the original fear memory and the extinction memory (4). While the current design was not able to optimally separate fear from extinction memory traces, future work could investigate whether extinction memories are prioritised for consolidation during SO-spindle coupling to enhance the strength of the extinction memory relative to the fear memory.

It has long been known that sleep disturbances are common in PTSD, and contribute to ongoing symptomatology (78). There has been growing interest in sleep spindles and their relevance to PTSD (27). Here, we contribute to this promising new area of research by examining the temporal coupling of spindles with slow oscillations, a mechanistic indicator of cross-regional dialogue and memory consolidation processes during sleep, in the context of PTSD. We found that PTSD patients, relative to trauma-exposed non-PTSD controls, exhibited delayed phase coupling, such that spindles coupled further away from the peak of the SO. Furthermore, the amount of coupling correlated with successful extinction retention, suggesting these oscillations to be critical in the successful consolidation of fear extinction.

## CRediT authorship contribution statement

**Dan Denis**: Conceptualization, Software, Formal analysis, Writing – Original draft, Writing – review & editing. **Ryan Bottary**: Conceptualization, Writing - review & editing. **Tony J. Cunningham**: Conceptualization, Writing – review & editing. **Per Davidson:** Conceptualization, Methodology, Writing – review & editing. **Cagri Yuksel:** Conceptualization, Methodology, Writing - review & editing. **Mohammed R. Milad** – Methodology, Writing – review and editing. **Edward F. Pace-Schott**: Funding acquisition, Conceptualization, Methodology, Supervision, Writing – review & editing.

## Data availability statement

The data analyzed in this study is subject to the following licenses/restrictions: the data underlying this article will become available in the future in the NIMH Data Archive (NDA) at https://nda.nih.gov, and can be accessed following instructions at https://nda.nih.gov/get/access-data.html. Requests to access these datasets should be directed to https://nda.nih.gov/get/access-data.html.

## Disclosures

This project was supported by NIMH grants R01MH109638 and R21MH115279 awarded to EP-S. Praxis Precision Medicines, Inc. provides partial salary support to EP-S. The remaining authors declare that the research was conducted in the absence of any commercial or financial relationships that could be construed as a potential conflict of interest.

## Supplementary Materials

### Skin conductance responses

SCRs were calculated for each trial as the mean skin conductance level in μS during the last two seconds of study context presentation subtracted from the maximum skin conductance level during the six seconds of colored lamp presentation. SCRs were square-root transformed, and recoded to zero in cases where the untransformed SCR was negative (1). Non-conditioning participants were excluded and were defined as those who exhibited less than 2 non-square-root transformed SCR responses to either of the two CS+s (in any combination) that were equal to or exceeding 0.05 μS during the Fear Conditioning phase (2). The first presentations of CS+1 and CS+2 during Fear Conditioning were excluded from analyses because their pairing with the unconditioned stimulus (shock) had not yet occurred. Similarly, these first presentations were not considered for the requirement of 2 non-transformed SCRs ≥ 0.05 μS.

### Extinction recall and generalization indices

The extinction retention index (ERI) was calculated as each participant’s average SCR to the first four CS+E trials of the extinction recall phase divided by their largest SCR to a CS+ trial during conditioning and multiplied by 100, yielding a percentage of the maximal conditioned fear that had been retained (2,3). Again, the first presentations of CS+1 and CS+2 during Fear Conditioning were excluded from use as the largest SCR to a CS+. Lower ERI indicates better extinction retention. The extinction generalisation index (EGI) was calculated by subtracting the average of the first four CS+E trials of extinction recall from the average of the first four CS+U trials of extinction recall, dividing this difference by the largest SCR to a CS+ trial during fear conditioning and multiplied by 100. This yields a percentage of the maximum conditioned fear represented by the retained difference in response to an unextinguished vs extinguished CS+ (4).

### Statistical analysis

Group differences in SCRs were examined via a linear mixed effects model with the factors Group (TEC, PTSD), Stimulus type (either CS+E, CS-for extinction recall, or CS+E, CS+U for extinction generalisation), Trial (first, second, third, fourth) and their interactions entered as fixed effects, and participant entered as a random effect.

## Results

### Behaviour

When comparing CS+E to CS-, there was a significant interaction between Stimulus type and Trial (*F* (1,614)=4.38, *p*=.036; **Figure S1**). On the first trial, SCRs to the CS+E were significantly higher compared to the CS- (*p*<.001). There was no difference between the CS+E and CS+U (*F* (1,624)=0.27, *p*=.61). No main effects or interactions involving group were observed (*p*s > .52).

### Associations between SO-spindle coupling and SCR measures of extinction

We ran exploratory regression analyses examining relationships between SO-spindle coupling and SCR-derived measures of extinction recall and generalisation. No significant associations were found (ps>.13).

**Figure S1.**
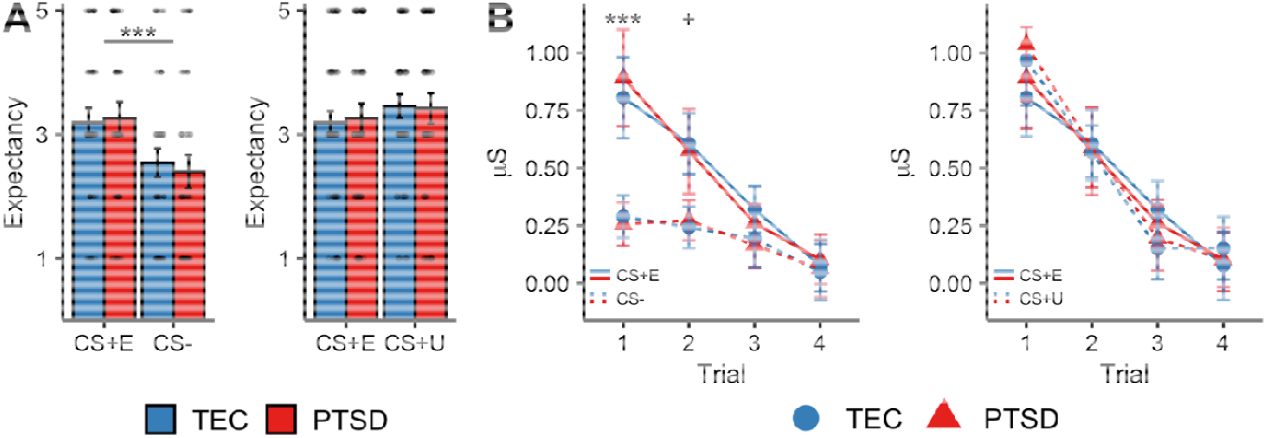
Behavioural and SCR results. **A** – Expectancy ratings to the first trial of each CS type. Expectancy ratings to the CS+E stimulus was significantly higher than the CS- (left panel), and no different to the CS+U (right panel). **B** – SCR responses to the first four trials of each CS type. Higher SCR responses were found to the first CS+E trial compared to the first CS- (left panel). No differences were found between CS+E and CS+U trials (right panel). *** = *p* < .001, + = *p* < .10. All error bars show the 95% confidence intervals around the mean.

